# Training Compute-Optimal Protein Language Models

**DOI:** 10.1101/2024.06.06.597716

**Authors:** Xingyi Cheng, Bo Chen, Pan Li, Jing Gong, Jie Tang, Le Song

## Abstract

We explore optimally training protein language models, an area of significant interest in biological research where guidance on best practices is limited. Most models are trained with extensive compute resources until performance gains plateau, focusing primarily on increasing model sizes rather than optimizing the efficient compute frontier that balances performance and compute budgets. Our investigation is grounded in a massive dataset consisting of 939 million protein sequences. We trained over 300 models ranging from 3.5 million to 10.7 billion parameters on 5 to 200 billion unique tokens, to investigate the relations between model sizes, training token numbers, and objectives. First, we observed the effect of diminishing returns for the Causal Language Model (CLM) and that of overfitting for the Masked Language Model (MLM) when repeating the commonly used Uniref database. To address this, we included metagenomic protein sequences in the training set to increase the diversity and avoid the plateau or overfitting effects. Second, we obtained the scaling laws of CLM and MLM on Transformer, tailored to the specific characteristics of protein sequence data. Third, we observe a transfer scaling phenomenon from CLM to MLM, further demonstrating the effectiveness of transfer through scaling behaviors based on estimated Effectively Transferred Tokens. Finally, to validate our scaling laws, we compare the large-scale versions of ESM-2 and PROGEN2 on downstream tasks, encompassing evaluations of protein generation as well as structure- and function-related tasks, all within less or equivalent pre-training compute budgets.

## 1 Introduction

Scaling up attention-based transformers has become a guiding principle for enhancing model performance across broad domains, particularly in Natural Language Processing (NLP) [12, 83, 67, 82, 4, 25, 82] and Computer Vision (CV) [94, 71, 21]. In recent years, large Transformer-based Protein Language Models (PLMs) such as PROGEN familiy [53, 63], ESM familiy [72, 49] and xTrimoPGLM [14] have also been developed, which leads to significant improvements over model performance on complex downstream tasks [47, 29]. Current pretrained language models utilize two main training objectives to encode sequence information: the BERT-like [24] Masked Language Model (MLM) and the GPT-like Causal Language Model (CLM) [11], each applied either separately or in a unified fashion. A common understanding is that bi-directionally encoded MLM excels in sample efficiency and shows enhanced performance in downstream task fine-tuning. This is particularly true in tasks that emphasize understanding complex patterns, making it a prevalent learning objective in modeling protein sequences [48, 14]. ^1^ On the other hand, uni-directional CLM, due to its sequential generation ability, is better suited for generating more coherent and realistic sequences compared to MLM [20, 63, 66].

However, training large protein language models (PLMs) are computationally intensive, and strategies for optimally allocating compute budgets for training PLMs are relatively underexplored, with *most efforts focusing on scaling model parameters based on a fixed set of training tokens to achieve performance improvements*. A key insight [44, 38, 79] is that large models should not be trained to their lowest possible loss to optimize computing; instead, models and data should be scaled proportionally based on available compute budgets. These scaling laws are broadly found in natural language models [44, 38, 35, 2, 60, 81, 18, 95]. But their applicability has not been validated within biological datasets, such as the primary structures of proteins, which are composed of amino acid sequences forming protein chains. Unlike natural languages, protein sequences are scientific data that are precisely represented using a vocabulary of 20 amino acids, with very little redundancy and are not as semantically smooth. Thus, we consider such data as a distinct modality and ask the question: *What are the scaling behaviors for MLM and CLM in protein language modeling?*

We focus on the best practices, which include revisiting datasets, optimization objectives, and model parameters as key factors. Our goal is to investigate an optimal training scheme for protein language models given predetermined compute budgets. Our core findings are as follows:

- We revisited the protein sequence data used for training PLMs and collected a dataset of 194 billion unique tokens on 939M unique sequences from publicly available sources to address the issue of overfitting and perform plateau in protein language modeling.
- We find that, in both MLM and CLM, training data scales sublinearly in the model sizes but follow distinct power-laws. MLM scales with a compute exponent of approximately 0.77. In other words, a 10*×* increase in compute leads to a 6*×* increase in MLM model size and a 70% increase in data, versus a 4*×* increase in CLM model size and a 3*×* increase in training tokens.
- We also find that models trained with CLM can be transferred to MLM. When given a predetermined amount of computation, and one wants to obtain both a CLM and a MLM model, there is a trade-off in allocating the training token to each model to jointly optimize the performance of the two. Interestingly, the allocation for CLM pre-training was determined by the scaling law of CLM and MLM, and the Effectively Transferred Tokens *D*_*t*_ from CLM to MLM. Furthermore, we verify this method experimentally using a 470M model and fine-tuning on downstream tasks.
- Building on our scaling strategies, we reevaluated the allocation of model size and training tokens under the compute budgets of established PROGEN2-xlarge and ESM-2 (3B) setups. Consequently, with the same compute budgets, we trained two corresponding models, one with 7.2B parameters and the other with 10.7B parameters, which exhibited enhanced performance in a diverse range of downstream tasks.

## 2 Scaling up data

First, we explore the effects of training PLMs across multiple epochs under token scarcity conditions. We then introduce a dataset, UniMeta200B, used throughout this work. This dataset enhancement alleviates the challenge of insufficient training for protein language models.

### 2.1 A Data-hungry Observation

Using the UniParc database with 250 million protein sequences, research on ESM [72] shows that the datasets UR50/S and UR50/D, with 45M and 65M unique sequences respectively, outperform Uniref100 in perplexity (PPL) on a ∼670M parameter MLM model. These datasets contain ∼15B and ∼20B unique amino acid tokens. The ESM-2 family models, ranging from 150M to 15B parameters, are trained extensively with nearly 1 trillion tokens over 45 epochs on the UR50/D dataset. In observing the scaling of ESM-2 models, it becomes apparent that increasing model size to 15B parameters from 3B shows marginal improvement. On the other hand, contemporary LLMs are often trained for only one or a few epochs [45, 38, 83, 84, 11]. The repetition of data with limited unique tokens has diminishing returns and hinders scaling model size [68, 36, 60, 74]. This underscores the importance of using rich datasets for training large-scale language models to ensure robust performance across applications.

We evaluated models with 150M and 3B parameters on the UR50/S dataset, trained on 200B tokens. The impact of token scarcity on PLMs’ scaling is shown in Figure 1, focusing on Independent and Identically Distributed (IID) validation and Out-Of-Distribution (OOD) test PPL, which measures the model’s randomness in amino acid selection. The smaller 150M model lacks good generalization, and increasing to a 3B model resulted in diminishing returns for CLM and severe overfitting for MLM, suggesting the CLM tasks are more challenging than MLM. In our settings, the output of CLM is computed by shifting the input position to the right for a single inference, ensuring the loss calculation encompasses all tokens in a sequence. The MLM approach masks 15% of the data during IID validation but sequentially masks each amino acid in a protein sequence during the OOD test. For our OOD dataset, we utilized the MMseqs2 tool [76] to conduct searches within the UniRef90 database for sequences post-training dataset timestamp, retaining those with *no detectable* identity. From these, a random sample of 3,000 sequences was selected to constitute the OOD dataset. Notably, we do not adopt dropout regularization, a practice that often reduces model capacity and is infrequently used in contemporary LLMs [45]. This choice is consistent with recent LLM configuration findings [40], including ESM-2 [49].

**Figure 1:**
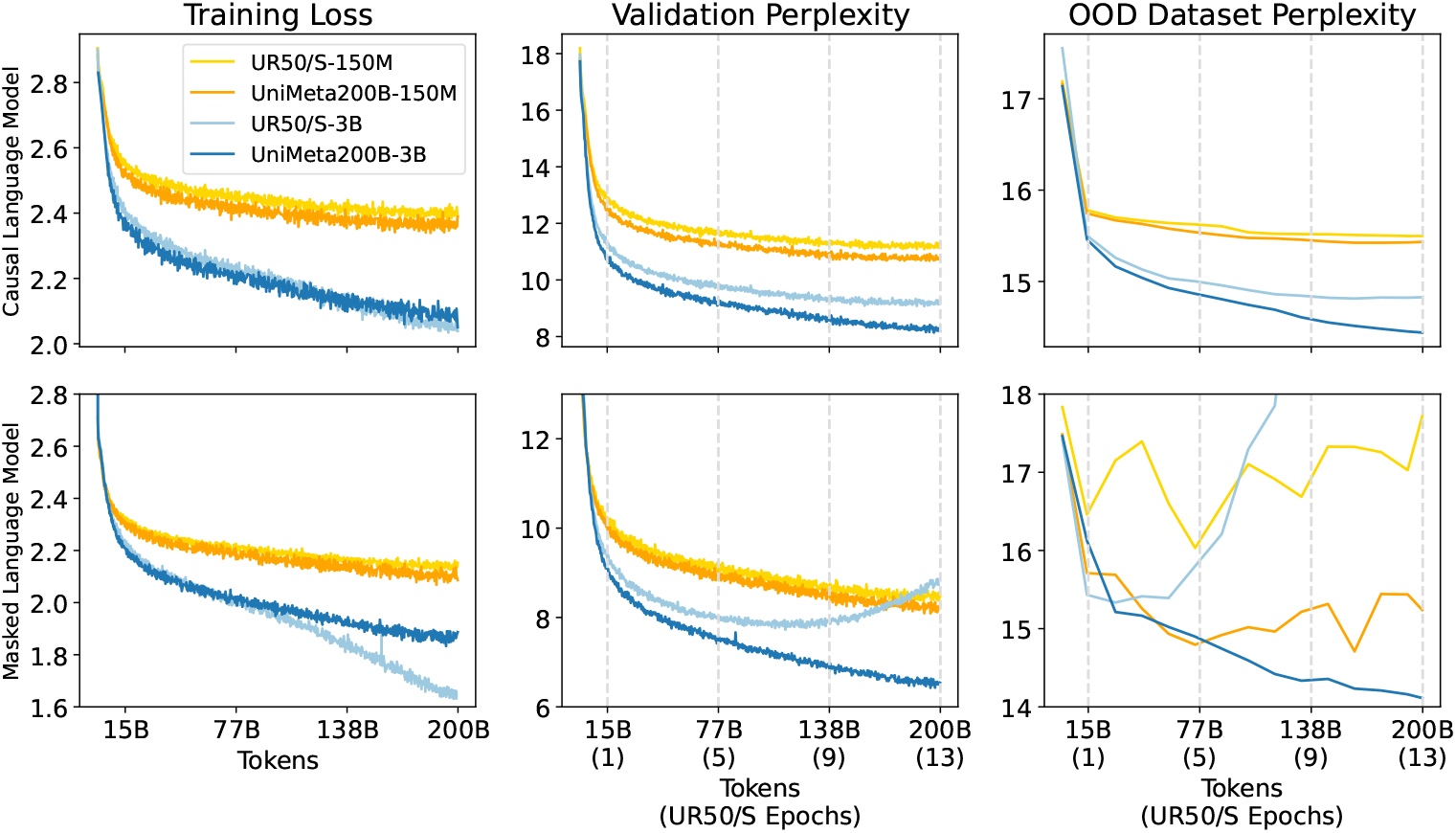
Learning curves for UR50/S and UniMeta200B. Training loss and validation PPL, OOD test PPL, were tracked over 200 billion training tokens for both the 150M and 3B models. As we scaled the model from 150M to 3B, we observed diminishing returns on CLM (First line) and a tendency to overfit on MLM (Second line) when repeating the Uniref50 (UR50/S) dataset. We totally evaluate 3 repeating methods on MLM 3B models, all of which present overfitting (see Appendix D).

### 2.2 Expanding Diversified Metagenomic Data

To tackle the challenge of data scarcity, we leveraged the Colab-FoldDB database [58], which focuses on metagenomic data sources such as BFD [1], MGnify [59], and specific eukaryotic and viral datasets including SMAG [23], MetaEuk [46], TOPAZ [3], MGV [61], and GPD [13]. We applied a stringent deduplication process with a maximum similarity threshold of 0.3 to preserve the diversity of the protein universe. Given that the Uniref90 dataset has proven most effective for pre-training across various Uniref clustering levels per ESM-1v [56], we incorporated Uniref90/50 (Before 2022-12), which includes incremental data relative to Uniref50/S representatives. ColabFoldDB_*c*_ and ColabFoldDB_*m*_ play dominant roles within the dataset, corresponding to cluster representatives and members, respectively. To ensure uniformity during training, we allocate weights within each batch to allow each amino acid token to be evenly processed through the model. This dataset, termed UniMeta200B, contains 939 million unique protein sequences and 194 billion amino acids, which is an order of magnitude larger than UR50/D. We observed significant improvements in the OOD test set and a consistent learning curve on the IID validation subset extracted from the training set (Figure 1). These enhancements not only ensure a controlled diversity to maintain sample efficiency but also significantly increase the quantity and uniformity of data, facilitating model scaling. ^2^

## 3 Parameters and Datasize Optimal Allocation

In this section, we propose a scaling law for protein sequences with MLM and CLM objectives, aiming at optimally balancing model size and data size under a fixed compute budget to improve efficiency on expanded resources.

### 3.1 Scaling laws for CLM and MLM

We first fit our models in the form of a fundamental power-law based on the existing work [44, 38, 35, 73, 2, 18, 95] in the field of LLMs. Specifically, given a fixed FLOPs formula of *C* = 6 *× N × D*, where *N* represents the number of forward-activated non-embedding parameters, and *D* is the number of training tokens, how should one navigate the trade-off between model size and the number of training tokens? The model parameters *N* and data size *D* can be directly fit with a simple power-law:

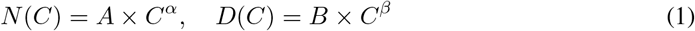

We employed the IsoFLOPs profiling approach [38, 9], setting 7 distinct training FLOP counts ranging from 1 *×* 10^18^ to 1 *×* 10^21^. For each FLOP count, we selected models from a pool of candidates (see Appendix O). Models were excluded if the estimated data size (*C/*(6 *\* N*)) resulted in more than 200B tokens or if the training steps were fewer than 20K. Ultimately, approximately 260 models were used for fitting. We considered the final validation loss for each model to ensure that every model completed a full cosine cycle with 10*×* learning rate decay. For each fixed FLOP count, we employ smoothed loss to determine the optimal model size with the smallest loss (Figure 2 (above)). Subsequently, we use Equation 1 and apply the least_squares method to fit the model.

**Figure 2:**
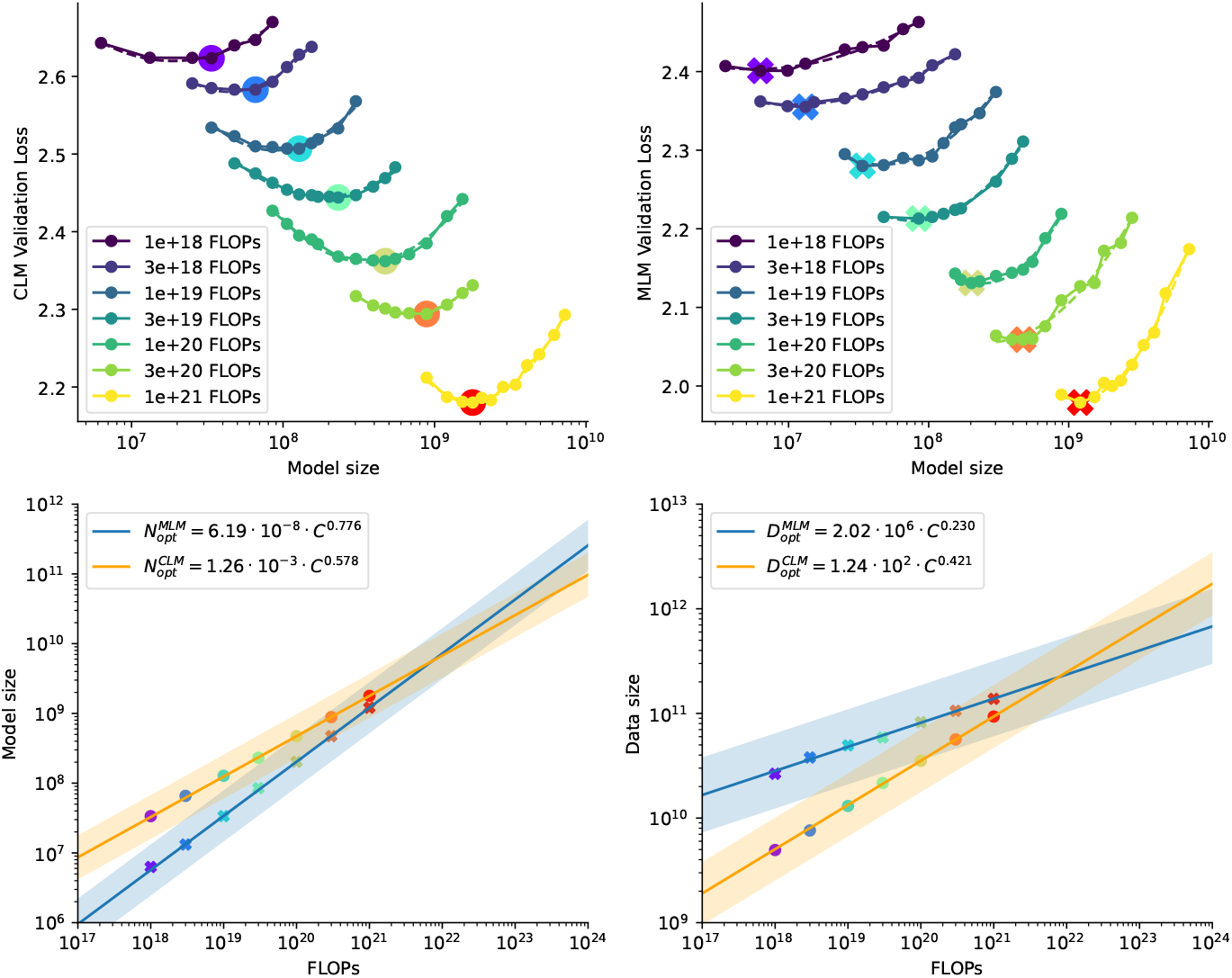
IsoFLOPs curves and parametric fit for CLM and MLM. We selected training tokens to ensure a uniform final FLOP count for different model sizes. The lowest loss of each curve revealed an optimal model size for a FLOP budget (**above**). We use these rainbow points at the valley to plot the efficient frontier for estimating the optimal model size and training tokens for scaling models (**below**). The interval range was estimated by model points with similar loss.

Given the minimal variations in the final loss among a set of (*N, D*) configurations, we classify these configurations as operating under “IsoLoss” conditions (see Appendix L Figure A15), considered optimal for training. In Figure 2 (below), we illustrate an efficient frontier interval that demonstrates permissible fluctuations in model size and dataset size at a specific FLOP count, while still achieving nearly identical losses. The variation in loss is quantified at 0.25 on a logarithmic scale with a base of 10. This indicates that within this FLOP counts, the model size can be adjusted within a range, increasing up to 80% or decreasing up to 40% without repeating data, to maintain a loss variation within 0.01.

We observe distinct growth rates in the proportional relationship between model size and training tokens for the MLM model compared to the CLM, as detailed in Table 2. Both models demonstrate an increase in the growth of model size that surpasses the growth of training tokens. Up to the intersection point around 1 *×* 10^22^ (see Figure 2, left below), the model size of MLM tends to be smaller than the CLM, thereafter, the MLM rapidly exceeds that of the CLM. Notably, the growth of the MLM’s training tokens is greatly lower than that for the CLM, possibly due to MLM’s higher sample efficiency. For instance, if the compute budget is increased by 10*×*, the size of the CLM model should increase by 4*×* and the training data by 3*×*, aligning more closely with equally proportional scaling. For the MLM, the model size should increase by 6*×* and the training data size by 1.7*×*.

**Table 1:**
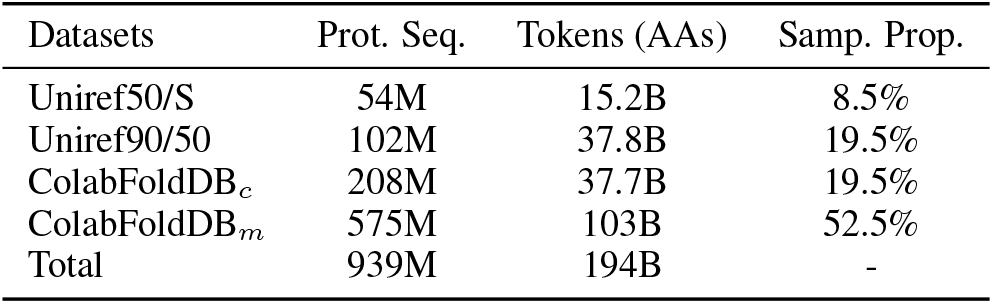
The Pre-training data. aggregates various public sources and specifies sampling proportions for a single epoch of training on 194 billion unique amino acids.

**Table 2:**
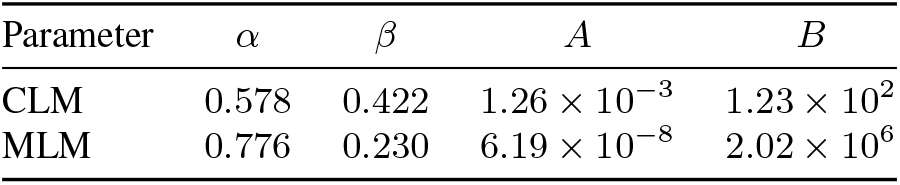
Coefficient of Equation 1.

In exploring the scaling relations of loss, we analyzed various model sizes *N*, compute budgets *C*, and training dataset tokens *D*. These can be described by a similar power-law relation defined as:

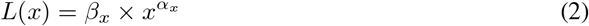

where *α*_*x*_ is the scaling exponent for different variables. For each FLOP count, we aimed to identify the minimal loss as the fitting target along with the corresponding independent variable *x*. Table 3 presents these fitting coefficients.

**Table 3:** Coefficient of Equation 2.

| Objective | $\alpha_N$ | $\alpha_D$ | $\alpha_C$ | $\beta_N$ | $\beta_D$ | $\beta_C$ |
| --- | --- | --- | --- | --- | --- | --- |
| CLM | -0.037 | -0.051 | -0.027 | 4.835 | 7.904 | 8.251 |
| MLM | -0.040 | -0.120 | -0.034 | 4.530 | 42.614 | 10.125 |

Based on the coefficients obtained from the fitting described above, we can establish the relationship between *D* and *N* by eliminating *L*. The relationship is expressed by the following equation:

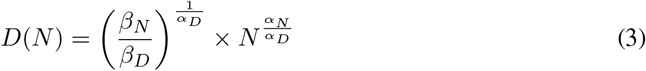

By substituting the learned coefficients into this formula, we can derive 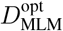 and 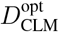 when given *N*. The estimation may be affected when the data exceeds 200 billion or when the quality or quantity of the training dataset changes.

### 3.2 Scaling law for training two models

When our goal is to optimize both CLM and MLM simultaneously, the strategic allocation of compute resources between these two objectives becomes essential. To facilitate this, we equalize model parameters across objectives to assess specific compute budgets for dual-objective training. Specifically, we seek the compute budgets, *C*_MLM_ and *C*_CLM_, for configurations where the optimal model size is the same, i.e., *N* (*C*_MLM_) = *N* (*C*_CLM_). These individual computations are then aggregated to formulate the overall compute budget:

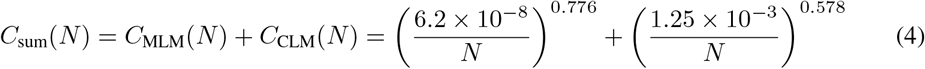

These two objectives share the same parameter size, their compute budget *C* and the number of training tokens *D* differ. Thus we further introduce a model-to-ratio *r*(*N*) as *D*_MLM_(*N*)*/D*_CLM_(*N*). We then achieve the relationship between *N* and *C*_sum_ by a fitted power-law (Figure 3) form:

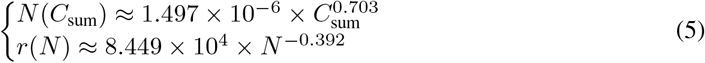

The ratio *r*(*N*) informs us about the allocation proportion of training tokens. Specifically, under equal parameters, the data for MLM should exceed that for CLM until a 10B threshold (achieving a 1:1) is reached, after which more training tokens are allocated to CLM.

**Figure 3:**
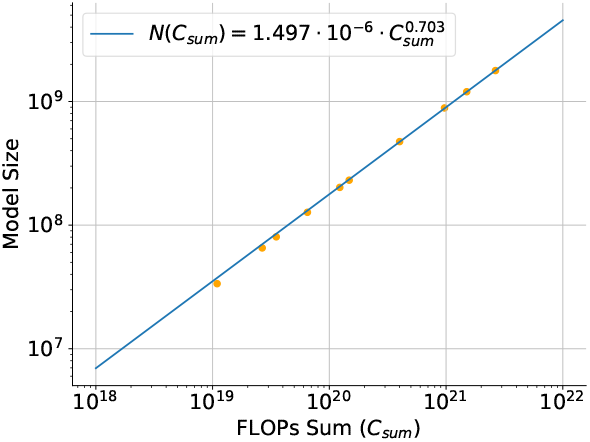
Compute allocation for two objectives with the same model size.

We further find that the scaling behavior of sparse parameter counts in a Mixture of Experts (MoE) model, set with eight experts (see Appendix J), as well as a combined power-law formula used to fit our data (see Appendix K), both exhibit a certain similarity to the scaling behavior we have proposed.

## 4 Transfer Scaling

We have outlined two independent scaling laws and how to allocate FLOPs under a fixed budget for training two optimal models, one with MLM and the other with CLM. However, we have not explored the interaction between these objectives. This raises important questions: Can models trained with one objective transferred to one with another objective? Is there a synergistic effect from training two models? Does training order impact the results?

### 4.1 Transferability

We conduct transfer learning experiments on MLM and CLM objectives, selecting eight optimal model sizes based on Equation 1. These models correspond to four increasing FLOP counts from 3 *×* 10^19^ to 1 *×* 10^21^ and undergo training from scratch followed by transfer training. Transfer training involves initially training on MLM or CLM, then training on the alternate model for each size.

We find that optimal pre-training on one objective benefits the target objective in transfer learning, though effects vary between methods. Starting with CLM and then training MLM, benefits increase with model scale. In contrast, starting with MLM then training CLM sees diminishing benefits. As shown in Figure 4 (left), for a model size of 230M with 3 *×* 10^19^ FLOPs, MLM from CLM pre-training reduces the loss by 0.02 compared to MLM from scratch, however, benefit that nears zero for the 1.7B model. Conversely, for models from 85M to 1.2B, transfer benefits grow with model size, the compared validation loss gap increasing from 0.025 to 0.045. This likely stems from the higher loss utilization rate in MLM; CLM calculates losses for all tokens in a protein sequence, whereas MLM only calculates losses for 15% of the tokens. ^3^.

**Figure 4:**
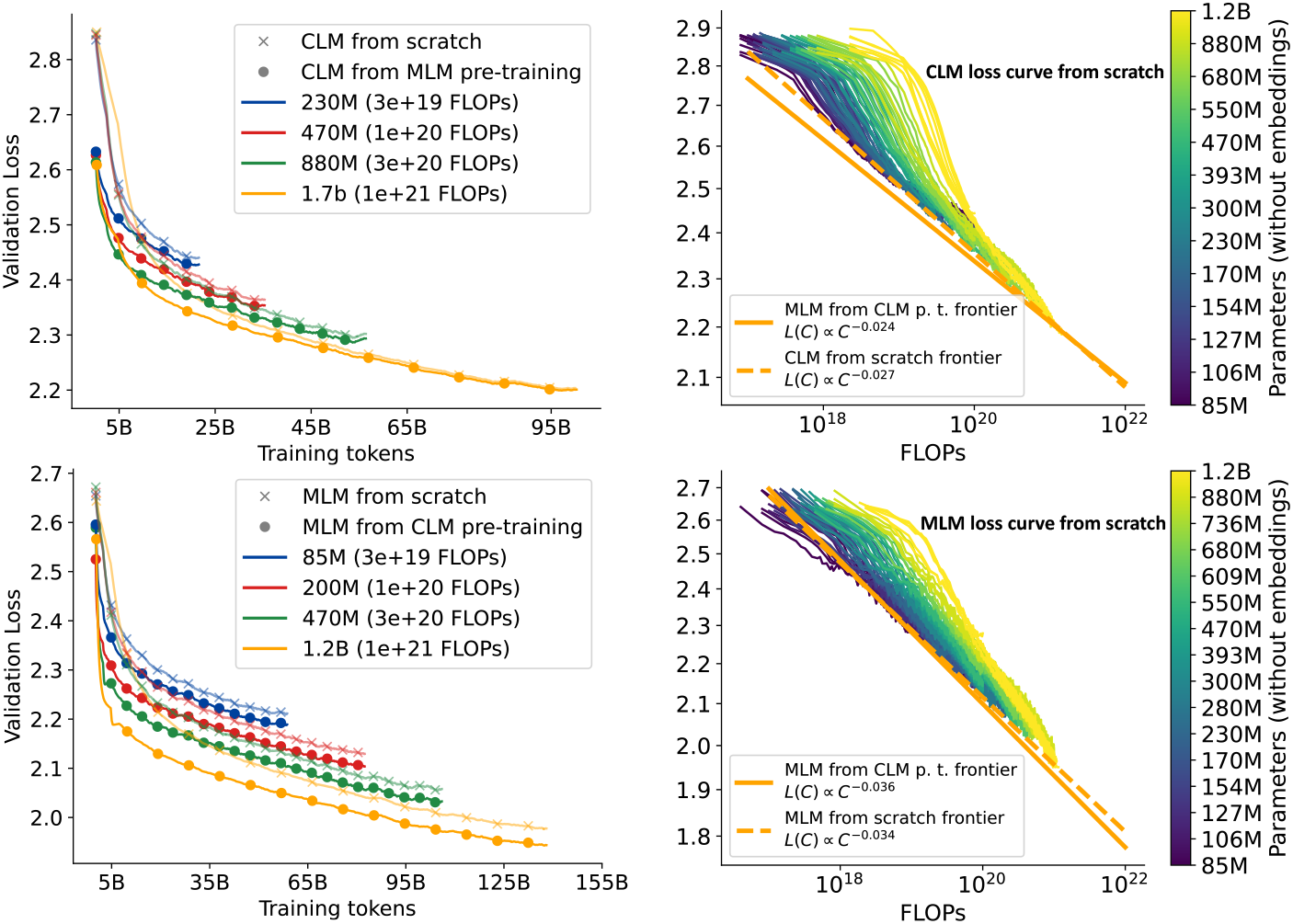
**Left:** The upper graph compares validation loss of CLM trained from scratch with those transferred from MLM, showing diminishing transfer benefits as model size increases. The lower graph depicts increased benefits for MLM from pre-trained CLM with larger sizes, indicating scale-dependent efficiency gains. **Right:** Shows loss curves for CLM and MLM across different FLOPs, emphasizing the efficient frontiers (or Pareto Frontier) from various transfer strategies. It highlights that the benefits of transferring from CLM to MLM grow with model size, reflecting a scale-dependent synergy between training objectives.

We use a power-law to model the transfer scaling law, initially excluding the pre-training FLOPs. The scaling behavior of transfer learning is modeled by:

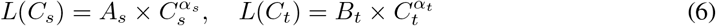

where *L*(*C*_*t*_) and *L*(*C*_*s*_) represent the loss for transfer learning and training from scratch.

Figure 5 (right) shows that the efficient frontier for *L*(*C*_*t*_) has shifted relative to *L*(*C*_*s*_) (it can be directly obtained from Table 3, repeated here for convenience.), indicating an improvement. The coefficients from both are shown in Table 4, where we can infer that 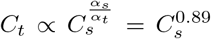, suggesting that training MLM from scratch with 10*×* the compute requires approximately 7.7*×* the compute compared to MLM from CLM pre-training. Another observation is that mixing training objectives in a single batch tends to be detrimental. Detailed results and settings are in Appendix I. The recommended transfer learning schedule involves pre-training CLM before MLM, as mixed training and order swapping show no significant benefits. We speculate that this primarily occurs because our MLM, which focuses solely on recovering corruption tokens, is not causal. If it predicted a middle segment in a left-to-right manner, it could mutually adapt with the context to accelerate training [90].

**Table 4:** Coefficients for *L*(*C*_*s*_) and *L*(*C*_*t*_)

| Parameter | $A_s$ | $\alpha_s$ | $B_t$ | $\alpha_t$ |
| --- | --- | --- | --- | --- |
| MLM | 10.125 | -0.034 | 11.133 | -0.038 |
| CLM | 8.251 | -0.027 | 7.191 | -0.024 |

**Figure 5:**
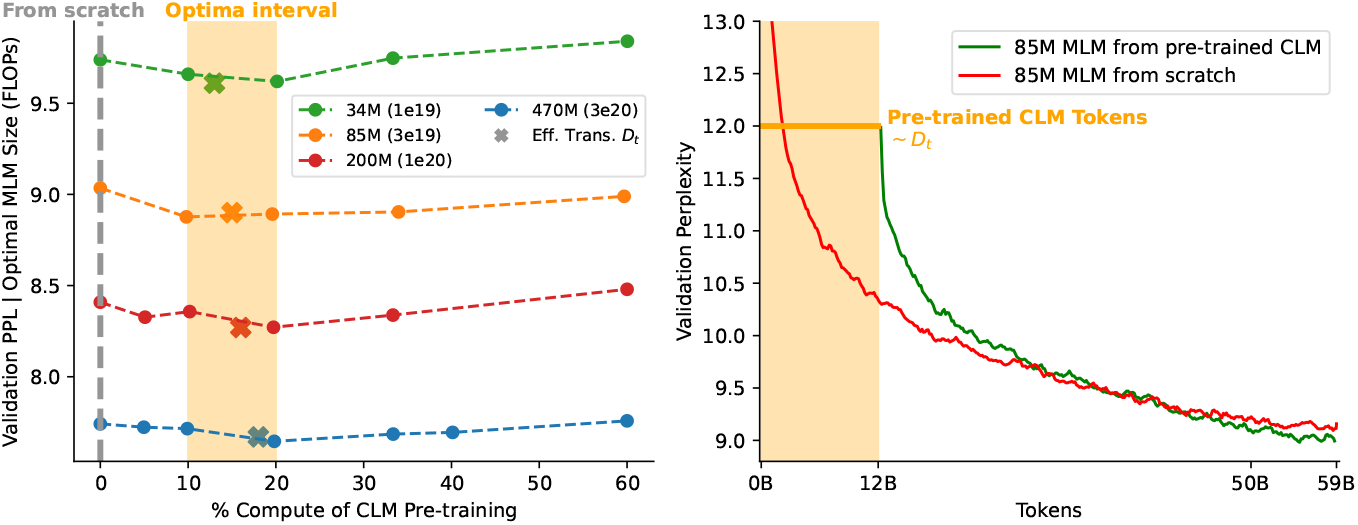
**Left**: Valid perplexity of % compute allocated for the CLM pre-training. For instance, % compute indicates first training on CLM and then the rest compute fine-tuning on MLM. The optimal CLM pre-training % compute range with [10, 20]. And the fitted *D*_*t*_*/*(*D*_*t*_ + *D*_*f*_) drops in the optimal loss range. **Right**: Comparison of validation perplexity for models trained from scratch (red) and those fine-tuned from a pre-trained CLM (green), demonstrating that fine-tuning from a CLM reduces perplexity with similar or even fewer tokens.

### 4.2 Effectively Transferred Tokens

Although we observe that MLM benefits from transfer learning from CLM, the pre-training compute budget remains unaccounted for. We focus on two aspects: (1) the actual benefit CLM provides to MLM and its predictability, and (2) performance differences between MLM trained from pre-trained CLM (MLM-CLM) and MLM from scratch (MLM-S) under identical FLOP constraints. We define Effectively Transferred Tokens *D*_*t*_ as the *additional data a model of the same size would need to train from scratch on MLM to achieve the same loss as a model pre-trained on CLM*. If the token number in the pre-trained CLM model exceeds *D*_*t*_, then the computations for CLM pre-training was excessive. Knowing *D*_*t*_ in advance would guide the allocation of tokens for CLM pre-training.

We compare MLM-S and MLM-CLM models ranging from 33M to 1.2B with FLOP counts from 3 *×* 10^19^ to 1 *×* 10^21^. By calculating the *token distance* at the same loss level between these models, we establish our fitting target *D*_*t*_, collecting approximately 2800 sample points. Following similar methods in scaling transfer works [35, 95], *D*_*t*_ is defined by a simple multiplicative scaling formula:

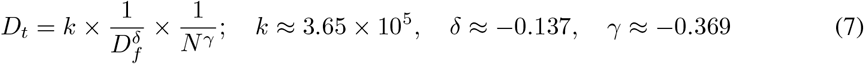

where *D*_*f*_ represents the tokens used for MLM-CLM, and *N* is the number of parameters, with *k, δ*, and *γ* as fitting coefficients. For instance, a 10*×* increase in *D*_*f*_ would roughly triple the model size and double *D*_*t*_. We validate these findings by evaluating the compute ratio of CLM pre-training under four specified parameters and FLOPs, as shown in Figure 5 (left), finding that MLM-CLM generally outperforms MLM-S. Specifically, *D*_*t*_*/*(*D*_*t*_ + *D*_*f*_) ranges from 10% to 20% of the compute budget for CLM pre-training. Figure 5 (right) schematically illustrates the learning curves of two 85M (3e19 FLOPs) models, with MLM-CLM achieving similar or better loss levels with equal or fewer tokens.

## 5 Experimental Validation

Based on the scaling laws we observe, we estimate the model size and training tokens for current leading models by analyzing their FLOPs. In our configuration, the PROGEN2-xlarge model, with 6.4B parameters, is estimated to require training with 7.2B parameters and 265B tokens. Similarly, the ESM-2 model, with 3B parameters, should be trained with a model size of 10.7B parameters and 260B tokens. Additionally, we employed two 470M models to test the transfer scaling strategy, one trained from scratch (470M scratch) and the other from CLM pre-training (470M trans.). The model’s details are reported in Table 5.

**Table 5:** Model architecture details. We compare popular models PROGEN2 and ESM-2 using similar FLOPs with our models estimated by proposed scaling law.

| Params | Objective | $N_{\text{head}}$ | Dim. | $N_{\text{layer}}$ | Train. Tokens | FLOPs |
| --- | --- | --- | --- | --- | --- | --- |
| PROGEN2-xlarge (6.4B) | CLM | 16 | 4096 | 32 | 350B | $1.34 \times 10^{22}$ |
| Our 7.2B | CLM | 32 | 4096 | 36 | 265B | $1.14 \times 10^{22}$ |
| ESM-2 (3B) | MLM | 40 | 2560 | 36 | 1T | $1.68 \times 10^{22}$ |
| Our 10.7B | MLM | 32 | 4352 | 47 | 260B | $1.68 \times 10^{22}$ |
| 470M scratch | MLM | 16 | 1280 | 24 | 106B | $3.0 \times 10^{20}$ |
| 470M Trans. | CLM + MLM | 16 | 1280 | 24 | 21B + 85B | $3.0 \times 10^{20}$ |

**Table 6:** Tasks performance of MLM Model on the test dataset with LoRA fine-tuning.

| Models | Contact Pred. (P@L/5) | Fold Class. (1195 class.) | Fluor. (reg.) |
| --- | --- | --- | --- |
| ESM-2 (3B) | 0.91 | 0.69 | 0.65 |
| Our 10.7B | 0.91 | **0.72** | **0.69** |
| 470M scratch | 0.78 | 0.65 | 0.67 |
| 470M trans. | **0.80** | **0.66** | 0.67 |

### 5.1 Protein Generation Comparison: 7.2B CLM vs. 6.4B PROGEN-xlarge

We first evaluate the perplexity on OOD data and then compare the protein generation capabilities of the 7.2B CLM and PROGEN2-xlarge models. Each model generated 2,000 sequences for each parameter combination of top-*p {*0.5, 0.7, 0.9, 1.0*}* and temperature *t {*0.2, 0.4, 0.6, 0.8, 1.0*}*, totaling 40,000 sequences per model. Sequences with a perplexity greater than 10 and duplicates were removed, leaving 8,263 and 8,466 sequences for the 7.2B CLM and PROGEN-xlarge, respectively. We used four metrics to assess the quality of the models and the generated sequences.

#### OOD Dataset PPL Analysis

We randomly sampled 5,000 sequences from UniProt released after 2023-01-01 and aligned them to our and PROGEN2’s training data (Uniref90 and BFD) using HHblits [70] or Jackhmmer [31]. Sequences below a maximum identity cutoff were used to assess the models’ PPL, as shown in Figure 6A. Our 7.2B CLM exhibited lower PPL on three subsets.

**Figure 6:**
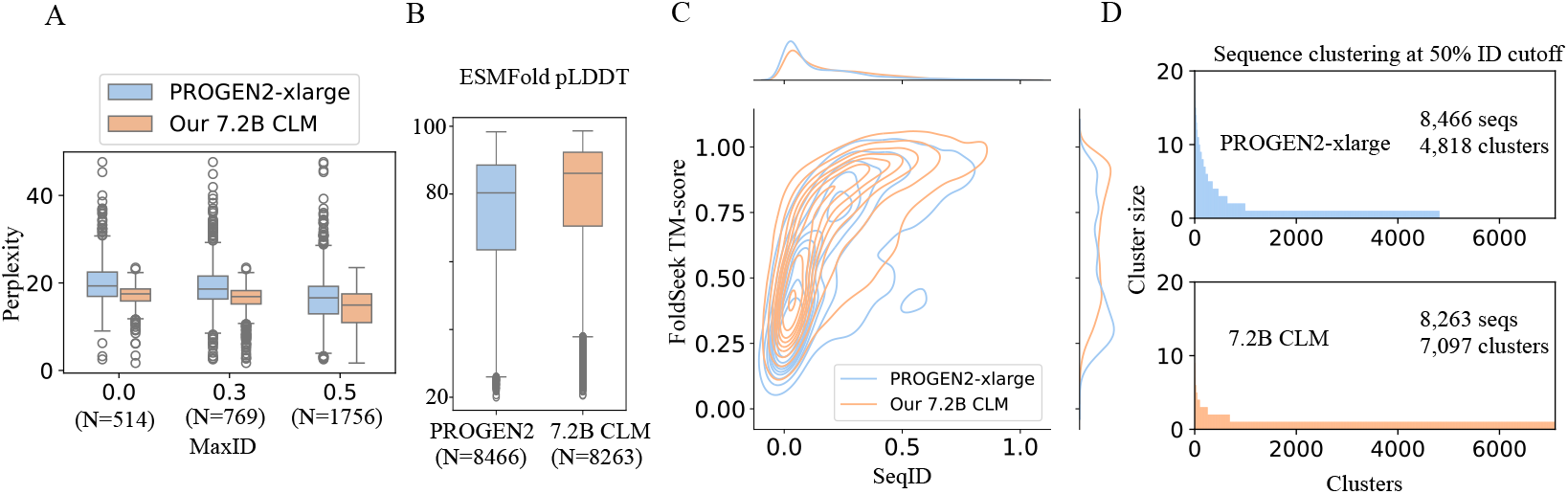
Comparative Analysis of CLM Models. **A**. Perplexity analysis for PROGEN2-xlarge and our 7.2B CLM shows lower values for our model across various MaxID levels, suggesting better sequence handling. **B**. Box plots of pLDDT scores for protein structures by PROGEN2-xlarge and our 7.2B CLM. **C**. Contour and line plots show our 7.2B CLM sequences mimic natural sequences more closely than PROGEN2-xlarge, assessed using Foldseek with the PDB database. **D**. Clustering at 50% sequence identity reveals our 7.2B CLM generates more clusters, indicating higher diversity. **Natural Sequences Comparisons with Foldseek** Using Foldseek [85], we searched the PDB database for sequences similar to those generated by our 7.2B CLM model, which showed better mimicry of natural sequence properties with higher average TM-scores (0.655 vs 0.522) and SeqID (0.194 vs 0.165), as shown in Figure 6C.

#### pLDDT scores from ESMFold

Atomic structures of 8,263 and 8,466 generated sequences were predicted using ESMFold, and compared based on pLDDT scores, displayed in Figure 6B. The 7.2B model’s average pLDDT score was 78.69, higher than PROGEN2-xlarge’s 74.33.

#### Diversity Analysis

Generated sequences were clustered using MMseqs2 [76] with a 50% similarity cutoff. The 7.2B CLM model resulted in higher diversity with 7,097 clusters compared to 4,818 clusters for PROGEN2-xlarge, detailed in Figure 6D.

### 5.2 Protein understanding tasks: 10.7B MLM vs. 3B ESM2

We evaluate task types: Contact prediction as binary classification at the amino acid pair level; fold classification into 1195 classes at the sequence level; and fluorescence as regression tasks. Following [14], we add a Multi-Layer Perceptron (MLP) head to each pre-trained model and apply Low-Rank Adaptation (LoRA) [39] (r=8, *α*=16) for fine-tuning (see Appendix H for convergence details).

#### Contact Prediction

This task determines if two residues, *i* and *j*, are in contact based on a distance threshold (<8Å). Uses the trRosetta dataset [28], split into 12,041 training, 1,505 validation, and 1,505 test samples [91]. Results are selected via validation performance. The evaluation metric is P@L/5 accuracy, considering residue pairs with a separation length greater than 6 and a sequence length cutoff of 512. We find a similar performance with ESM-2 (3B) and interestingly, larger ESM-2 (15B) is even slightly worse than ESM-2 (3B), with the results being closely matched overall at 0.91. On the other hand, the 470M model from CLM pretraining (470M trans.) outperforms the model trained from scratch (470M scratch).

#### Fold Classification

This task assigns protein sequences to one of 1,195 known folds, primarily identifying novel remote homologs. This task is significant in proteomics and structural biology for analyzing folding patterns and advancing disease research [15]. It shows improved accuracy over ESM-2 (3B), which we expect is due to increased data diversity scaling. Additionally, the 470M model transferred from CLM pre-training continues to perform effectively in this task.

#### Fluorescence

The fluorescence task predicts the fluorescence intensity of green fluorescent protein mutants. Following the TAPE splitting method [69], dataset sizes are 21.4K for training, 5.4K for validation, and 27.2K for testing. The evaluation metric is the Spearman score. Our 10.7B model, shows promising results on this task, while the performance difference between the two 470M models is minimal.

## A Conclusion

We first expanded metagenomic databases such as ColabFoldDB, emphasizing the critical importance of data quality and quantity for scaling language models. Then we introduced scaling laws for two distinct language model tasks and provided recommendations on the allocation of model size and data size under expanding compute budget, along with the potential loss prediction. Additionally, we explored the transferability between these tasks. Our findings were validated by robust performance across various downstream tasks. We expect that they will extend to and assist other domains and biological data modalities.

## B Related Work

### Protein Language Model

Since the advent of AlphaFold2 [43], the masked language model (MLM) has been integrated as a subtask within the Evoformer architecture. In this context, an assumption is that large language models can be considered as a lossless compression method [22]. This was followed by a series of language modeling efforts [33, 10, 34, 30, 29], which aimed to conduct pre-training on single-sequence proteins using larger datasets and model scales. These efforts sought to harness the scale of the models to learn complex co-evolutionary information, although detailed investigations on how to optimally scale these models remain scarce. Our work primarily focuses on these finer aspects, aiming to fill this gap in the research.

### Training objectives

In natural language processing (NLP), masked language models (MLM) are rarely adopted due to the self-explanatory nature of natural language, which inherently prompts the meta-knowledge of tasks and generates task targets through CLM (Conditional Language Modeling) training models. However, a unified language modeling objective for Protein Language Models has yet to be fully consented. Those based on causal language modeling (CLM) have been primarily explored for protein design. Benchmarks in protein design using MLM [88] have also shown promising results for generation [64], exhibiting variable performance when compared to CLM [96, 87]. Additionally, the potential of the in-filling task objective remains largely unexplored [6, 80, 26]. Our research aims to thoroughly discern the scaling behavior of the two most common optimization objectives in this domain.

### Scaling Laws

To our knowledge, the concept of scaling laws of language model is first introduced by OpenAI [44]. Subsequently, numerous variants and modifications [38] have been developed around this theme. Recently, an array of new scaling laws has emerged. These include scaling laws related to learning rates and batch sizes [9], data-constrained scaling laws [60], scaling laws for downstream tasks and Transfer [95, 37], as well as scaling laws within the Mixture of Experts (MoE) framework [18], and those concerning long sequences and positional encoding [51]. While these laws are primarily derived using auto-regressive models in resource-rich domains, their application in the biological data sector is less common. Our work seeks to address this gap. Furthermore, scaling laws for Masked Language Models (MLM) are notably scarce. Given that MLMs are currently one of the most effective training methods for biological data, our research on MLMs could also be extended to other non-text domains.

## C Discussion and Limitations

### Repetition for Masked Language Model

Our scaling law is learned within a single epoch setting. It is well known that MLM exhibits higher sample efficiency than CLM due to the variable masking of training samples across epochs. However, this advantage diminishes when training is limited to only one epoch. This also suggests that for MLM training, a small amount of repetition can be considered as new data, without detriment to the performance. We present empirical evidence comparing a 2.8B model trained on 1T tokens (approximately five epochs) against a 10.7B model trained on 265B tokens (roughly 1.4 epochs). Despite the models utilizing the same amount of FLOPs, the latter achieves optimal training by attaining lower out-of-distribution (OOD) perplexity (10.33 vs 10.21). Despite this, the impact of training MLM for several epochs repeatedly is not significant in terms of loss. This insight suggests that repeating several rounds under MLM training has a minimal impact on reducing loss, and our scaling law does not necessarily need to be confined within 200 billion tokens. And smaller models are more user-friendly during inference and fine-tuning. Therefore, we also suggest an alternative approach that adjusts the optimal training token count and model size within our scaling law framework when scaling MLM. We will further investigate repeat scaling laws as designated in future work [60].

### Multi-modality Scaling

We observe that the scaling laws for CLM, also known as autoregressive models, exhibit similarities to those in natural languages or the code modality in the context of protein sequences, closely aligning with findings by Chinchilla [38]. The multi-modal auto-regressive work [35] suggests the existence of a nearly universal scaling law across various modalities, including images, videos, math, code, and languages. Our results appear in this trend as well. The same situation may apply to other modalities of biological data, such as RNA and DNA [62]. However, for the scaling laws pertaining to MLM, we have yet to identify detailed research on or evidence of a universal scaling law. This gap highlights a crucial area for future investigations, potentially extending our understanding of scaling across different model architectures and data modalities.

### Hyperparameters Sensitivity

When selecting language models and configuring their training processes, model size and number of training tokens are not the only hyperparameters that require careful selection. Other critical factors, such as the learning rate schedule, batch size, also play significant roles. We rely on existing work and provided experimental heuristics to determine the other necessary hyperparameters [93, 84]. The maximal learning rate (LR) was determined using a LR finder [65] to prevent model collapse, then empirically choosing the median value where the loss decline is steepest. Our observations suggest that while the exact maximum learning rate was not highly sensitive, ensuring a steep decline in loss and the completion of the learning rate schedule was crucial, typically with an error loss of around 0.01. The critical batch size [55] and model loss are correlated [44], indicating that larger models require correspondingly larger batch sizes; we empirically set models under 1B parameters to a batch size of 512K and models over 1B to 1 million.

### Other Dataset and Strategies

While our datasets encompass a significant portion of the protein universe, they might still not be fully representative. The combination of BFD [8], Uniref [78], MetaClust [46], and IMG/JGI [54] with 90% clustering includes at least 600B unique tokens. However, different datasets might induce variations in the power-law behavior, such as changes in the slope or shifts in the log-log space, which warrants further investigation. It may be of interest for future work to test the applicability of our findings to different model architectures. Currently, there is significant research into the scaling of LLMs for long sequences [17, 16, 7, 92, 19, 50, 41], and the MSA augmentation could notably enhance protein representation in terms of contacts and structure. Exploring the scaling laws in this context may be an interesting avenue for future work.

## D UR50/S Repeat Experiments

We employed three different methods to repeat training on the UR50/S dataset, all of which ultimately led to overfitting. The reference for these experiments is shown by the blue curve in Figure A7, which represents UniMeta’s loss for approximately one epoch.

Firstly, using bootstrapping, we processed 200 billion tokens from UR50/S with replacement. In each epoch, 65% of the dataset was randomly selected, leading to a diminished proportion of unsampled tokens by the fifth epoch, as depicted by the orange curve.

Secondly, we shuffled the unique data for each epoch to ensure that all UR50/S tokens were used per epoch, resulting in a stair-step pattern [32] in the training loss, illustrated by the green curve. It has simply memorized the dataset but isn’t improving at generalizing. Over-confident predictions of the first batch of the next epoch lead to a big step update, and then the model is not adapted to the next batches, resulting in no longer a decrease in loss.

Lastly, we shuffled the entire training dataset less stringently, which did not strictly ensure that all UR50/S tokens were used every epoch, but guaranteed that each token was used an equal number of times over the entire training period. We term it global shuffle, this approach is shown by the red curve.

From the gradient norm curve shown in Figure A7 (right), we observe an uptick in gradient norm for the overfitting curves, indicating that the model is no longer optimizing effectively. In machine learning, such an increase in gradient norm typically suggests that the model is encountering areas of the parameter space where gradients are steeper or more erratic, often occurring when the model starts to memorize the training data rather than generalize from it, approaching a saturated network [57]. This behavior can result from overly complex models, too many training epochs without sufficient regularization, or training on non-representative data.

**Figure A7:**
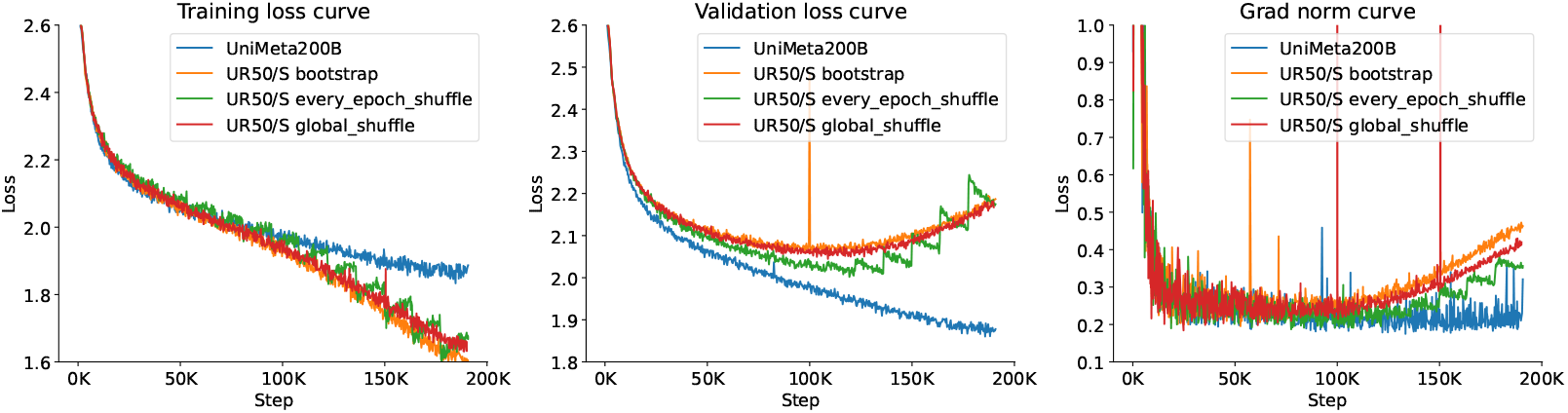
Learning curve for UR50/S dataset repetition methods. Our 194B tokens dataset (UniMeta200B) shown in blue, serves as the reference with an approximate single epoch run. The bootstrapping method, depicted in orange, processes 200 billion tokens with replacement, indicating a tendency towards zero unsampled tokens by the fifth epoch. The every-epoch shuffle method, in green, ensures all tokens are used per epoch, forming a stair-step pattern in training loss. Lastly, the global shuffle method, in red, loosely uses all tokens each epoch but ensures the strict number of epoch passes for every token. The rightmost plot of gradient norms shows an uptick for curves corresponding to overfitting, signifying a lack of further optimization, with steep or erratic gradients indicated by the ascending gradient norms.

## E Choice of Masking Ratio

**Figure A8:**
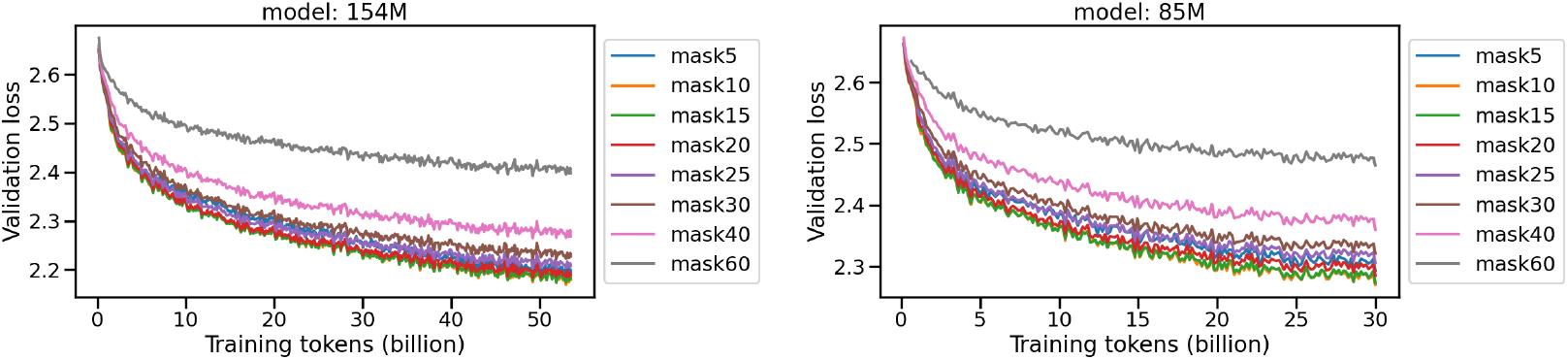
Validation loss of different masking ratios. Two models (154M and 85M) are trained from 5% to 60% masking intervals.

In the original BERT work [24], the absence of masked tokens in downstream tasks presented a mismatch with the pre-training data distribution. The authors investigated various masking ratios and concluded that a 15% masking rate was most beneficial for downstream tasks. This was implemented alongside an 80-10-10 strategy: 80% of the tokens were replaced with a mask, 10% were randomly substituted, and the remaining 10% were left unchanged.

However, given the significant differences between protein sequences and natural language processing data, we employed two models, sized at 85M and 154M, to explore a range of masking ratios from 5% to 60% (see Figure A8). The best masking ratios for validation loss drop ranged from 10% to 20%; ratios too small (5%) or too large (greater than 25%) degraded the performance.

We further used pre-trained eight different models to perform full fine-tuning on downstream tasks such as Contact Prediction and Fold Classification in Figure A9. Results from the test datasets revealed that, similar to NLP, the optimal performance was achieved within a 10%-20% masking range. Specifically, a 20% masking ratio slightly outperformed 15% in Contact Prediction, while the 15% ratio yielded the best results in Fold Prediction. Consequently, for our Masked Language Model (MLM), we decided to adhere to the 15% masking ratio with the 80-10-10 strategy for training all our models.

**Figure A9:**
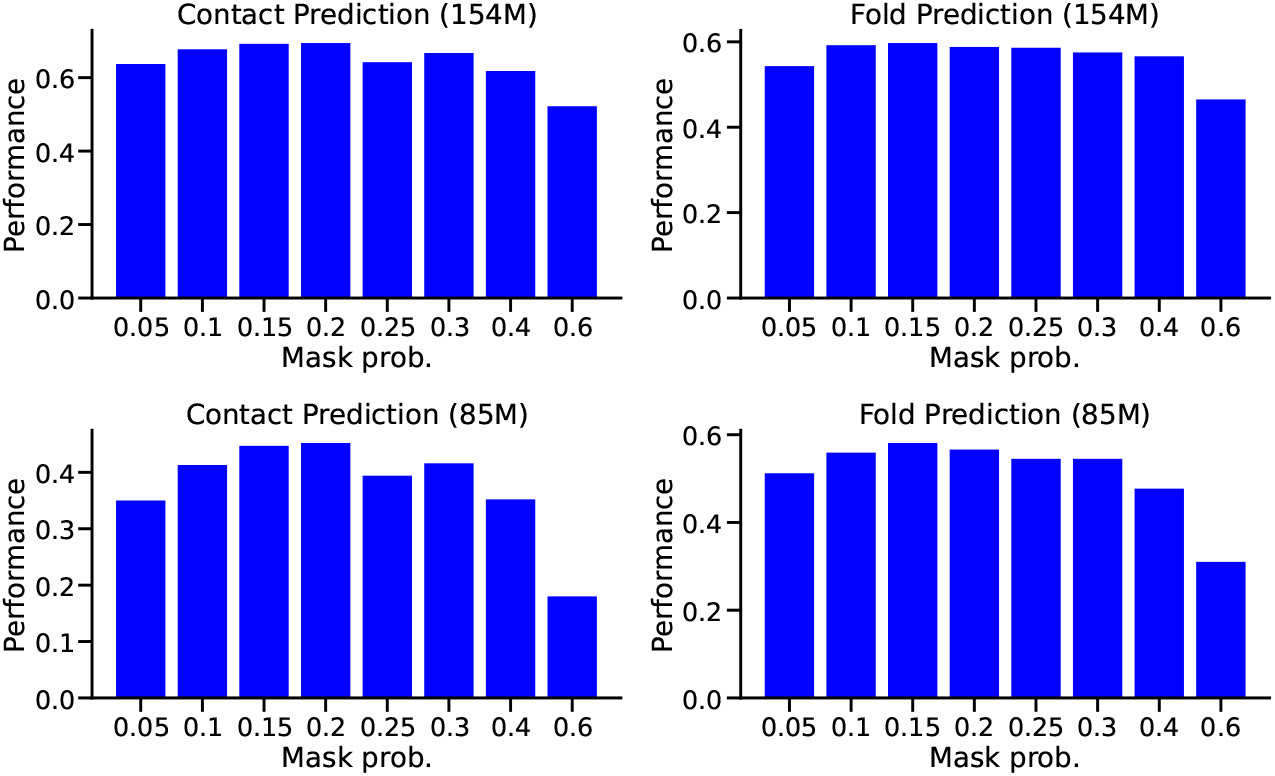
Abalation of different masking ratios. Two models (154M and 85M) are trained from 5% to 60% masking intervals, and evaluated on contact map and fold classification downstream tasks.

## F MLM/CLM for Protein Contact Prediction

**Figure A10:**
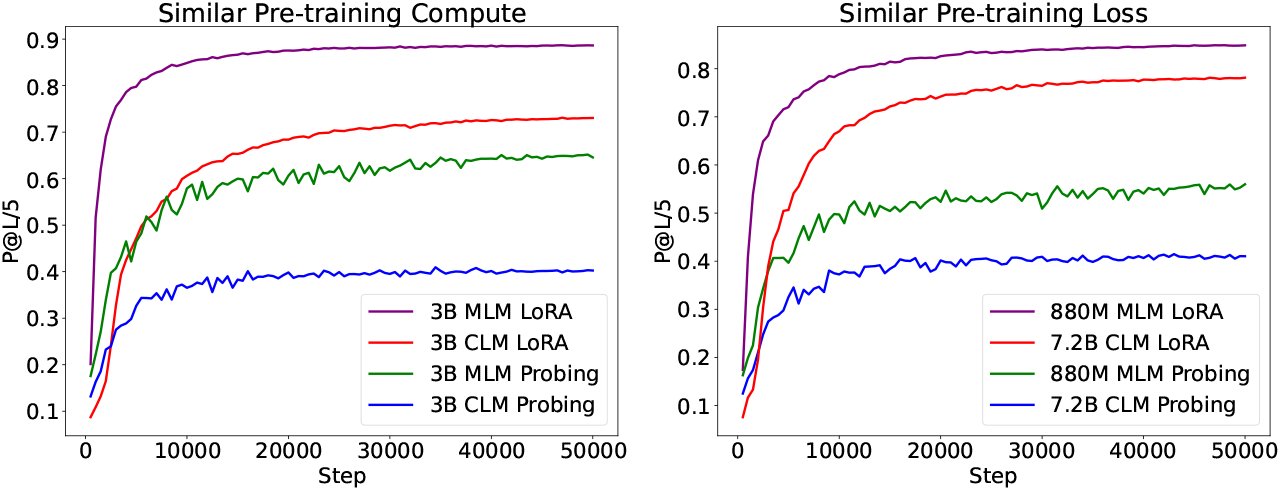
Contact Prediction on MLM and CLM models. Two 3B models (CLM and MLM) were trained using identical computational resources, represented by the probing and LoRA fine-tuning methods. On the right, performance of a 7.2B CLM model is compared with an 880M MLM model under similar pre-training loss conditions. These models exhibit differing rates of convergence, highlighting the impact of uni-directional and bi-directional model architectures on learning dynamics.

We compared the effectiveness of CLM in the downstream task of contact prediction, using two different setups (Figure A10). In the first setup, two 3B models were trained under identical computational resources on 200 billion tokens, 3.4 *×* 10^21^*FLOPs*. Their performance was evaluated through two training approaches: Probing (freezing the pre-trained model) and LoRA fine-tuning, with an added MLP head for comparison.

In the second setup, we compared the effects of MLM and CLM under similar loss conditions. Here, a 7.2B CLM model and an 880M MLM model were selected, both achieving a loss of 1.98 on our validation set. Despite the MLM model having a simpler loss calculation, involving a 15% mask rather than a one-by-one mask—which would result in a higher loss—the MLM significantly outperformed the CLM. Importantly, the CLM model’s computational power was an order of magnitude greater than the MLM model (1.68 *×* 10^22^ vs 1.0 *×* 10^21^ FLOPs). This suggests that despite the lower loss achievable by the CLM model compared to MLM with a one-by-one mask, the unidirectional limitations of CLM do not translate into better downstream task performance.

## G Pre-training Dataset Quality

Compared to Uniref90, ColabFoldDB offers a higher diversity and larger numbers of protein sequences, though with generally shorter sequence lengths, likely suggesting potentially lower data quality. To evaluate the efficacy of our expanded dataset, ColabFoldDB, we initially trained two 85M models separately on Uniref90 and ColabFoldDB. Uniref90 in our dataset comprises two subsets: Uniref50/S and the incremental dataset over Uniref50/S, termed Uniref90/50. Similarly, ColabFoldDB consists of representative and member data. We controlled the sampling proportion to ensure uniform sampling across both datasets, with results reported in Table A7. Both models were then trained using identical configurations on a 50B scale.

From the perspective of validation loss in pre-training, the higher loss on ColabFoldDB might be attributed to its lower diversity and shorter sequence lengths compared to Uniref90. However, the performance on downstream tasks, such as contact prediction and fold classification, shows negligible differences between models trained solely on ColabFoldDB and those trained on Uniref90, as illustrated in Figure G. This confirms that ColabFoldDB is an effective expansion of Uniref90 that maintains sample efficiency.

**Table A7:** Compared two dataset characteristics. Protein sequence count, token number, and sampling proportions for Uniref50/S, Uniref90/50, and ColabFoldDB representative and member data.

| Datasets | Prot. Seq. | Tokens (AAs) | Sampling Prop. |
| --- | --- | --- | --- |
| Uniref50/S | 54M | 15.2B | 28.67% |
| Uniref90/50 | 102M | 37.8B | 71.33% |
| ColabFoldDB<sub>c</sub> | 208M | 37.7B | 26.75% |
| ColabFoldDB<sub>m</sub> | 575M | 103B | 73.52% |

**Figure A11:**
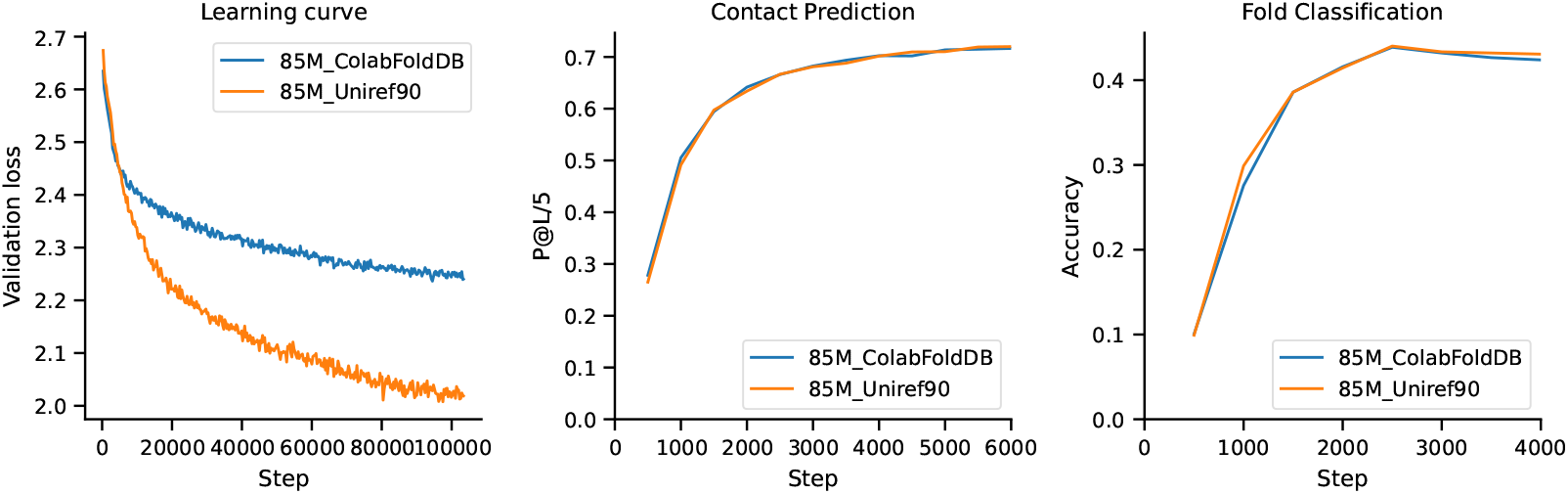
Data quality check. Comparison of learning dynamics and downstream task performance for two 85M models trained on ColabFoldDB and Uniref90. Left: Validation loss curves demonstrating initial training differences. Middle: Contact prediction performance showing the response to testing on similar tasks. Right: Fold classification accuracy, comparing model responses to structural prediction tasks. Despite initial differences in loss, both datasets yield comparable performance in downstream applications.

## H Convergence of Downstream Fine-tuning Tasks

Observing the learning curves in Figure A12a, we can assess the effectiveness of different fine-tuning scenarios. For the contact prediction task, the convergence speed under the LoRA setting is very similar for both models. Our testing reveals closely matching results for ESM-2 models with capacities of 650M, 3B, 15B, consistent with the findings reported by Ankh et al. [29]. This similarity suggests possible saturation of the dataset under single-sequence pre-trained models. Additionally, the convergence rates for tasks such as fold classification and fluorescence are generally faster than those for ESM-2, indicating robust generalization following our data augmentation strategies.

Based on the two 470M models defined in our Table 5, despite using the same computational power, we observe distinct outcomes (Figure A12b) in contact prediction and fold classification tasks. The MLM model from CLM pre-training converges slightly faster than MLM from scratch. However, the distinction is less pronounced in the two downstream regression tasks. This suggests that perplexity is more sensitive to protein structure related tasks, i.e., contact prediction and fold classification, but shows less sensitivity to regression tasks, particularly when assessed using the Spearman metric, which is prone to variability.

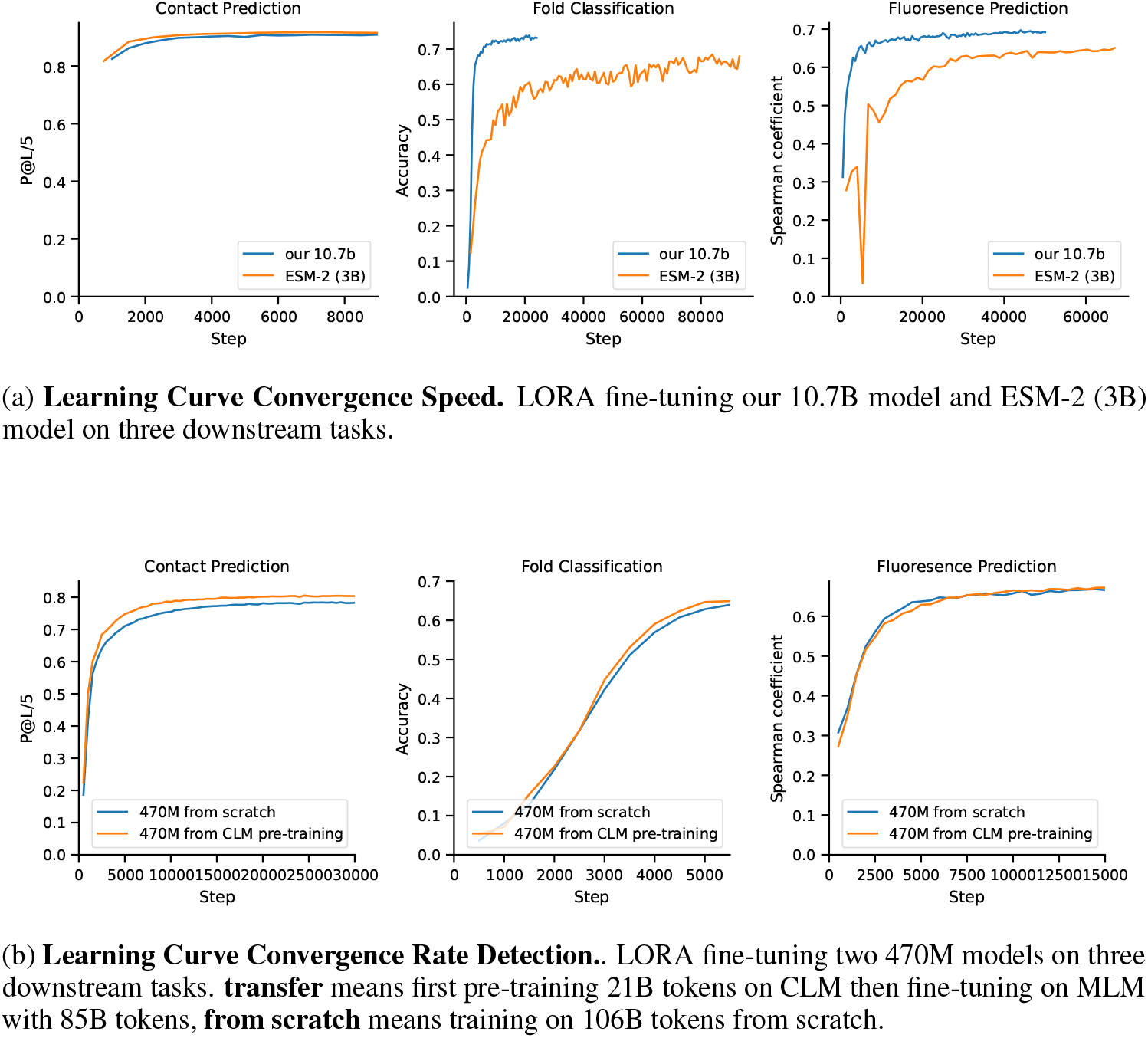

## I Mixed Objectives Training

We also employed an untied model to simultaneously optimize two objectives:

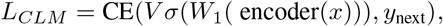

and

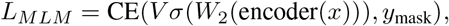

where *V* represents the protein vocabulary embedding, and *W*_1_ and *W*_2_ are the parameters corresponding to the CLM and MLM tasks, respectively. CE is the cross-entropy operator. The *σ* is the Tanh activation function.

We compared CLM and MLM under our scaling law of optimal model and data size distributions. One approach involved training from scratch, while the other used mixed training. In the mixed training approach, the actual number of training tokens was higher due to the additional FLOPs consumed by another optimally trained objective, in other words. In other words, mixed training consumes the FLOPs of two optimal allocations; we only extracted the loss curve of one target for comparison. We extracted the loss curve of just one target for comparison with the from-scratch training. Our findings indicate that mixed training of the two targets can lead to detrimental interference, an effect not observable in smaller models, as depicted in Figure A13. The possible reason for this situation is that mixed training has reduced the batch size for one of the objectives, making optimization difficult. We did not further investigate the impact of increasing the batch size and only observed based on the training tokens. However, we cannot rule out the possibility that they are mutually detrimental. Therefore, if both objectives are to be optimized concurrently, a sequential training strategy should be employed: first optimizing CLM, followed by MLM training. We consider that CLM is more challenging to predict than MLM, which may allow the model to capture more complex and implicit sequential features initially, thereby enhancing its ability to understand and predict masked words in subsequent MLM training.

**Figure A13:**
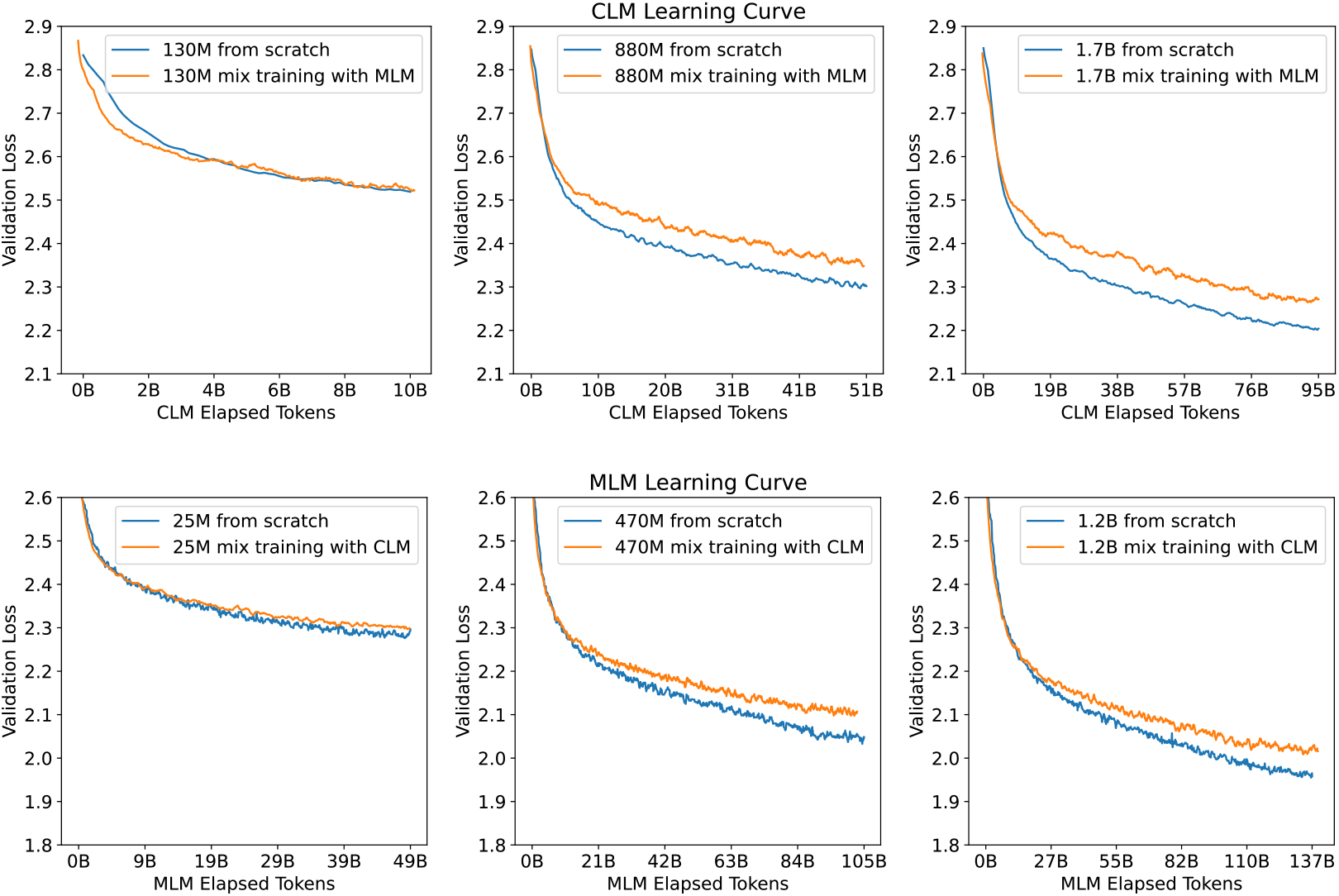
Mixed objective validation loss. Comparative validation loss curves for models trained from scratch versus mixed training approaches. Each panel corresponds to different model sizes, as indicated by the parameters. For each model, two training strategies were compared over an identical number of elapsed tokens: training from scratch (blue) and mixed training with the other objective (orange). Across all model sizes, training from scratch consistently achieves lower validation loss compared to mixed training, suggesting that mixed training may not be as effective as dedicated training for each individual objective.

## J MoE Scaling

We find that the scaling behaviors of sparse parameter counts in Mixture of Expert (MoE) models are remarkably similar to those of dense model sizes, potentially allowing for a reduced compute budget for modeling scaling behaviors due to less activated parameters per token.

In our experiments, we evaluate MoE models ranging from 10M to 500M sparse parameter counts, using a model size of 17 with eight experts, following the settings outlined in Mixtral of experts [42], including its load-balancing scheme. The figure below shows different IsoFLOPs curves. Notably, the FLOPs here are calculated based on sparse parameters rather than actually activated ones. We use the method described in the main text to select optimal loss points and fit these around the sample points, enabling us to project the optimal model size and number of tokens for larger models (center and right). We observe that the power-law coefficients for CLM and MLM are similar to those of dense models, with MoE CLM vs. Dense CLM at approximately 0.57 vs. 0.58, and MoE MLM vs. Dense MLM at 0.74 vs. 0.77.

**Figure A14:**
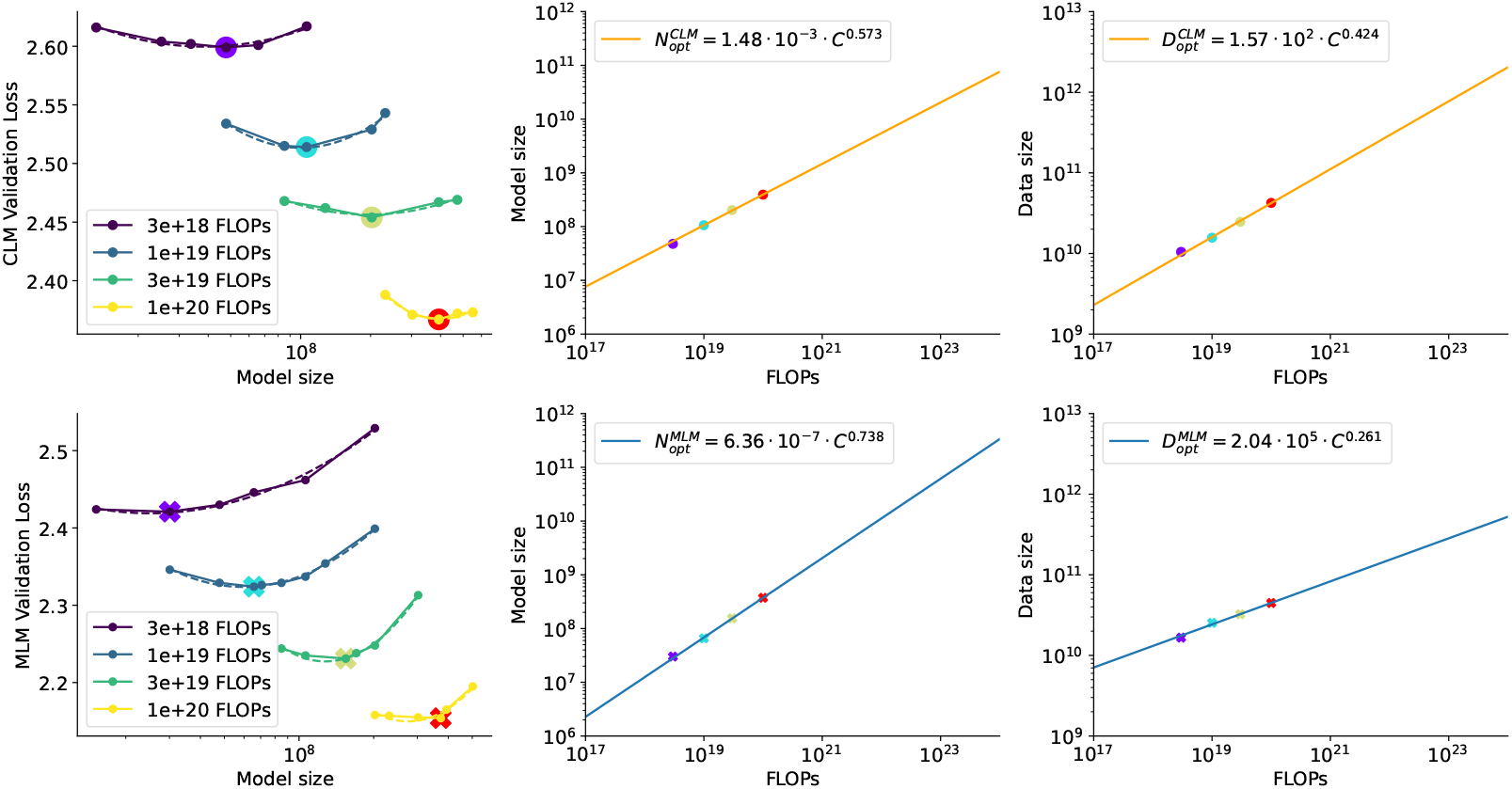
Scaling laws of MoE. The scaling behaviors of sparse parameter counts (8 experts) in MoE models, highlighting IsoFLOPs curves for different model sizes and FLOPs configurations. Each graph represents the relationship between model size, FLOPs, and validation loss for both CLM and MLM using MoE configurations. The power-law fits indicate optimal model size and data requirements for efficient scaling, showing that MoE models closely align with dense models in terms of scaling efficiency, with power-law coefficients for MoE-CLM and MoE-MLM approximating those of their dense counterparts. This suggests that MoE models can achieve similar scaling behaviors with potentially lower computational costs.

Our study strictly focuses on models with eight experts, which may not be entirely rigorous. Clark et al. [18] proposed a unified scaling law defining effective training parameters for MoE, aiming to harmonize the scaling laws for Dense and MoE models. The investigation of biological data will be considered in future work.

## K Combined Power-law

We applied the fitting function proposed by Chinchilla [38], detailed in Equation 8, to model the effects of various factors on model performance. It can provide a loss prediction where neither the parameters or model size are not optimal allocation. This loss function simultaneously depends on parameters *N* and *D*:

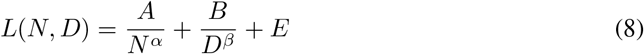

where *E* denotes the irreducible loss. Parameters *A, B, α*, and *β* are learned through the fitting process. As *N* → ∞ or *D* → ∞, the function degenerates to a form similar to Equation 2, which indicates that it models the scenarios under perfect conditions of other variables.

Given that most of our training tokens are used for less than or equal to one epoch, and that the model size is prone to underfitting at fixed FLOPs, the asymptotic behaviors *L*(*N*) at *D* → ∞ and *L*(*D*) at *N* → ∞ are enough for determining the parameters in *L*(*N, D*).

To enrich data points, we randomly added several FLOP counts into 25% of the model size and trained these models for 0.25, 0.5, 0.75, and 1 epoch. And we adopt the Huber loss to fit these coefficients:

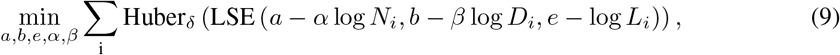

where LSE represents the log-sum-exp operator, and *δ* = 10^−3^. The terms *N*_*i*_, *D*_*i*_, and *L*_*i*_ denote the model size, dataset size, and loss of the *i*-th run, respectively. We fitted the MLM validation loss from 110 samples and the CLM validation loss from 149 samples using grid search with *α* ∈ *{*0, 0.5, …, 2*}, β* ∈ *{*0, 0.5, …, 2*}, e* ∈ *{*−1, −0.5, …, 1*}, a* ∈ *{*0, 5, …, 25*}*, and *b* ∈ *{*0, 5, …, 25*}*. The final initialized parameters of CLM and MLM both are [*e, a, b, α, β*] = [1, 5, 10, 0.5, 0.5]. We set the maximum number of iterations to 1000, and the two objectives were essentially achieved after 360 iterations. The exponential powers of learned *a* and *b* yielded the coefficients *A, B*, which were reported in Table A8.

**Table A8:** Coefficient of Equation 8

| Table A8: Coefficient of Equation 8 |  |  |  |  |
| --- | --- | --- | --- | --- |
| Objective | $A$ | $B$ | $\alpha$ | $\beta$ |
| CLM | 143.9 | 22036.5 | 0.367 | 0.496 |
| MLM | 3.365 | 7.569 | 0.042 | 0.099 |

Substituting all learned coefficients into the following Equation from the original Chinchilla paper:

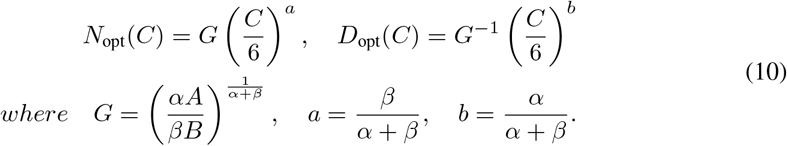

The results closely approximate the trends given in Equations 1 and 2, confirming our overall findings.

## L IsoLoss

In addition to using the seven different FLOPs counts reported in the main text to determine the optimal model sizes and fit our scaling law, we also incorporated additional model points into our analysis. We trained using the final loss points of all the CLM and MLM that are run. Figure A15 depicts the contour of the fitted function *L* and the efficient frontier as a red dashed line, presented in log-log space. The frontier interval of Figure 2 is computed from this observation. From this approach, it revealed the scaling exponents for model size to be 0.77 in MLM and 0.57 in CLM, very similar to the IsoFLOPs profiling method in Section 3.1.

**Figure A15:**
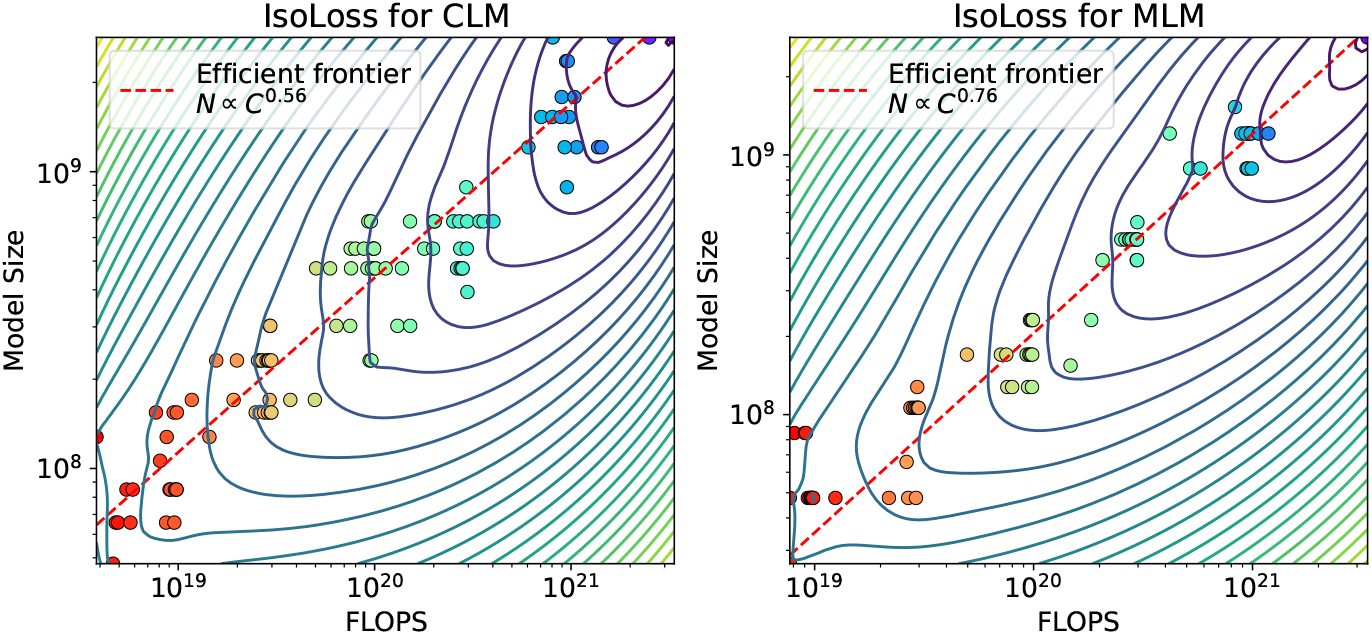
Parametric fit for CLM and MLM. Unlike the IsoFLOPs method used in the main text to select the optimal model size, these plots use all available data points to fit the models. The left panel shows the contour of the function *L* and the efficient frontier (indicated by the red dashed line) for the CLM, and the right panel for the MLM. The rainbow dots represent identical loss. The results closely align with using the IsoFLOPs profiling method.

## M Training Procedure

We conducted all experiments using Ampere A100 GPUs (80G) equipped with NVLink, utilizing the GLM framework [93, 27] developed based on DeepSpeed and Megatron. We have used a total of around 1 million GPU hours. Our approach predominantly utilized data parallelism, avoiding model parallelism and pipeline parallelism to simplify deployment. Modifications were made to the standard Transformer architecture [86], adopting a DeepNorm [89] strategy and layer normalization [5]. The activation function was set to GLU [75], RoPE [77] was used to encode position, similar to the settings found in the Transformer++ architecture [83]. We further adopt FlashAttention [19] to accelerate our training process. The used max LR empirically found to range between 6 *×* 10^−4^ and 1.2 *×* 10^−4^ from small to large model size, was used along with a cosine decay strategy to reduce it to 0.1 *×* max LR. Both CLM and MLM were trained under similar settings for model size, with a consistent LR and a minimum warm-up period of 2.5% steps, extending to at least 100K training steps. All sequences were set to a length of 1024, with sequences concatenated using an <EOS> delimiter. Based on findings related to loss magnitude and batch size [55]. The AdamW optimizer [52] was used with *β*_1_ = 0.9, *β*_2_ = 0.95, *ϵ* = 1 *×* 10^−8^, and a weight decay of 0.01. All experiments omitted the dropout (it reduced the capacity to hinder model scaling) and trained with bfloat16. Most pre-training experiments were confined to the ≤ 1 epoch, with some models extending up to 30% beyond one epoch. For the transfer learning setting, we load the finished checkpoint of the pre-training model and disregard the pre-trained optimized state, and learn rest tokens with warmup 5% steps the max LR.

## N Broader Impact

If the scaling law of the protein language model improves predictions or understanding of protein structure and function, it could potentially have positive impacts on scientific research in fields such as biology, medicine, and drug development. This may facilitate the development of new drugs, accelerate progress in disease diagnosis, or drive advancements in frontier research in the life sciences.

## O Model Parameters

Table A9 details the sizes and configurations of all models utilized in this research, training only with data parallel expcept 10B with tensor parallel size 2:

**Table A9:**
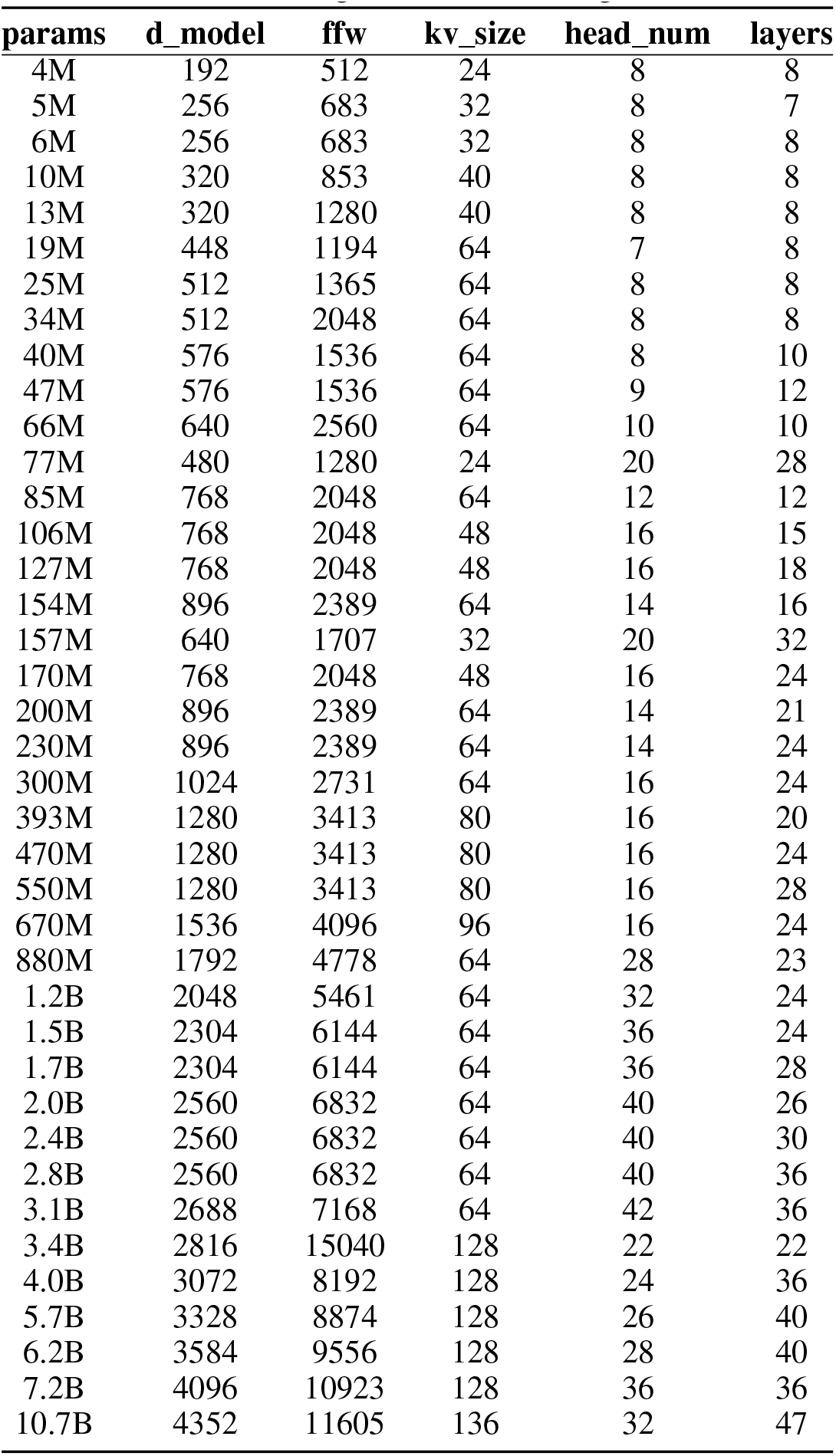
All model hyperparameters. Several of the models presented have been trained using various learning rate schedules and differing amounts of training tokens.

## Footnotes

1 Appendix F also compared CLM and MLM on the protein contact prediction task through fine-tuning and freeze probing, with MLM demonstrating superior performance relative to CLM.

2 Appendix G compare the training performed separately on two datasets, and we find that the ColabFoldDB does not affect downstream results.

3 Appendix E analyzes the mask ratios.

## Notes

### Competing Interest Statement

The authors have declared no competing interest.

